# DNA barcoding of fungal specimens using long-read high-throughput sequencing

**DOI:** 10.1101/2022.02.08.479507

**Authors:** Kadri Runnel, Kessy Abarenkov, Ovidiu Copoț, Vladimir Mikryukov, Urmas Kõljalg, Irja Saar, Leho Tedersoo

## Abstract

Molecular methods are increasingly used to identify species that lack conspicuous macro- or micromorphological characters. Taxonomic and ecological research teams barcode large numbers of collected voucher specimens annually. In this study we assessed the efficiency of long-read high throughput sequencing (HTS) as opposed to the traditionally used Sanger method for taxonomic identification of multiple vouchered fungal specimens, and providing reference information about intra-individual allele polymorphism. We developed a workflow based on a test-set of 423 fungal specimens (representing 205 species), PacBio HTS method, and ribosomal rRNA operon internal transcribed spacer (ITS) and 28S rRNA gene (LSU) markers. PacBio HTS had a higher success rate than Sanger sequencing at a comparable cost. Species identification based on PacBio reads was usually straightforward, because the dominant operational taxonomic unit (OTU) typically represented the targeted organism. Unlike the Sanger method, PacBio HTS enabled detecting widespread allele polymorphism within the ITS marker in the studied specimens. We conclude that multiplex DNA barcoding of the fungal ITS and LSU markers using a PacBio HTS is a useful tool for taxonomic identification of large amounts of collected voucher specimens at competitive price. Furthermore, PacBio HTS accurately recovers various alleles, which can provide crucial information for species delimitation and population-level studies.

## Introduction

A large proportion of living organisms can reliably be identified to species only based on molecular methods. Collected and vouchered specimens constitute a permanent record of biological diversity and form the basis of taxonomy as they provide material for description of new species and reference for taxonomic identification. The modern species identification relies increasingly on informative marker genes – termed as DNA barcodes (Hebert et al., 2003) – and sometimes on the entire genomes of organisms (Coissac et al., 2016; Misas et al., 2020). Because the DNA of non-living specimens degrades rapidly (Taylor & Swann, 1994), it is feasible to retrieve the molecular information in a reasonable time frame (usually within a few years following collection) to prevent loss of valuable genetic information. Large research teams of ecologists and taxonomists typically collect hundreds to tens of thousands of specimens annually, which necessitates efficient bulk analysis of large amounts of specimens in terms of cost, time, and labor (Hebert et al., 2018; Srivathsan et al., 2018).

DNA barcoding for taxonomic identification is traditionally performed using the Sanger sequencing method. Typically, one or several marker gene fragments are amplified and sequenced, allowing to produce up to 1000 base pair high-quality reads in a single pass, depending on the purity of DNA. The Sanger sequencing technology relies on chain termination signal averaged across all amplicons, and produces a consensus read of sequences of several marker gene alleles from the target specimen (Sanger et al., 1977). A well-known limitation of this approach is that the consensus read blurs the evolutionary signal among alleles. Additional technical shortfalls include low quality in the beginning of the sequence, disruption of reads in the case of length polymorphism in alleles, generation of ambiguous signal in the case of nucleotide substitutions, and failure to produce a readable sequence if both the target and co-occurring organisms (e.g. contaminants) are amplified or when the amplicon is of low quantity or purity (Hyde et al., 2013).

To overcome the issues with Sanger sequencing, high-throughput sequencing (HTS) methods can alternatively be used for DNA barcoding (Coissac et al., 2016, Bohmann et al., 2020). These methods include HTS-based analysis of single or multiple marker genes, genome skimming (i.e., low-coverage genome sequencing to retrieve long contigs of marker genes), and whole-genome sequencing. Genome-based methods are useful for obtaining full-length marker genes for accurate identification and phylogenetic analyses, but they have a low capacity to phase alleles differing by a few substitutions or indels >500 bases apart (Coissac et al., 2016; Tedersoo et al., 2016). In addition, such methods are relatively costly, because each sample requires preparation of a specific library and assembly from hundreds of thousands to tens of millions of reads. Furthermore, no more than one or two marker genes are still broadly used for DNA barcoding, except multi-gene phylogenetic analyses. Short-read HTS platforms such as Illumina, Ion Torrent and DNBseq provide high-quality reads for DNA fragments <550 base pairs, which may be insufficient for reliable identification and phylogenetic analyses. In comparison, long-read and synthetic long-read sequencing methods offer great promise for analysis of DNA markers up to ca. 3500 base pairs (Hebert et al., 2018; Callahan et al., 2021; Karst et al., 2021). Despite high raw error rate, these methods are highly accurate when calculating consensus (built-in option for PacBio and synthetic long-reads). Long-read methods also enable to phase haplotypes, which is of great relevance in population-level research (Tedersoo et al., 2021). Although these methods are relatively costly, hundreds to thousands of samples can be multiplexed for a single run, bringing the overall costs comparable to Sanger sequencing and one to three orders of magnitude less compared with genome-based approaches (Hebert et al., 2018; Srivathsan et al., 2018).

The main purpose of this study was to assess the efficiency of long-read HTS for taxonomic identification of large sets of vouchered fungal specimens and providing reference information about intra-individual allele polymorphism. The latter is important to avoid describing artefactual “ shadow taxa” (i.e., based on sequencing artefacts, rare alleles and pseudogenes; Thines et al., 2018; Porter & Hajibabaei, 2021) and inflating HTS-based biodiversity estimates. Here we developed a workflow based on PacBio multiplex DNA barcoding method and a test set of hundreds of fungal amplicons using the ribosomal rRNA operon internal transcribed spacer (ITS) and 28S rRNA gene (LSU) markers. These are the two most commonly used DNA barcodes for taxonomic identification of fungi and many protist groups (Pawlowski et al., 2012; Schoch et al., 2012). We demonstrate that the benefits of this multiplex DNA barcoding method include better recovery of low-quality samples and higher read quality compared with Sanger sequencing, as well as retrieval of multiple alleles differing by a single or more base pairs at a comparable cost.

## Materials and methods

### Molecular analyses

To test the relative performance of Sanger and PacBio sequencing on fruiting body samples, we compiled 423 vouchered specimens (polyporoid, resupinate, and agaricoid fruiting body types) belonging to 205 species from the fungarium of Natural History Museum of Tartu University (acronym TUF; Appendix 1). Most fruiting body samples were collected between 2015 and 2020 (Table S1) for different ecological and taxonomic studies.

The DNA of specimens was extracted from roughly 0.1-1 mg dried material using ammonium sulphate lysis buffer (Anslan & Tedersoo, 2015) that provides sufficient DNA quality from fruiting body material (Tedersoo et al., 2016). For all samples, our DNA barcoding approach targeted the ITS region, and for one batch of samples we also analyzed the LSU. To cover the entire ITS region, we chose the primers ITS1catta (5’-ACCWGCGGARGGATCATTA-3’) and ITS4ngsUni (5’-CCTSCSCTTANTDATATGC-3’) for PCR. The ITS1catta primer has a high affinity to Dikarya and it avoids amplification of the common intron in the 3’ end of the 18S region (Tedersoo & Anslan, 2019). Both primers were tagged with one of the 115, 12-base, sample-specific indices (Tedersoo & Anslan, 2019). PCR was carried out in two replicates in the following thermocycling conditions: an initial 15 min at 95 °C, followed by 30 cycles of 95 °C for 30 s, 55 °C for 30 s, 72 °C for 1 min, and a final cycle of 10 min at 72 °C. PCR products from replicate samples were pooled and their relative quantity was estimated by running 5 µl DNA on 1% agarose gel for 25 min. DNA samples with no visible bands were re-amplified with 35 cycles. To retrieve the LSU, we also amplified the DNA from a subset of 75 samples using the untagged primer LROR (5’-ACCCGCTGAACTTAAGC-3’) and tagged primer LR5 (5’-TCCTGAGGGAAACTTCG-3’) and the above-described options. In total, our test set yielded 497 fungal amplicons.

The PCR products were checked on a 1% agarose gel and the relative strength of the band was recorded at the scale of 0 (no band) to 5 (very strong band). It was also recorded whether there were a single or multiple bands on the gel. Altogether 20 µl of the PCR products were subjected to purification using Exo-SAP enzymatic treatment (Tedersoo et al., 2006) and shipped for Sanger sequencing in Macrogen, Inc., the Netherlands, using a single-pass with the untagged ITS4ngsUni primer (or LROR primer for LSU). Of the remaining PCR products, between 1 and 10 µl of amplicon were taken based on the strength of the band on the gel (categories 0 and 1, 10 µl; category 5, 1 µl), and pooled into five sequencing libraries for the ITS region and one library for the LSU region. The amplicon pools were purified using FavorPrep™ Gel/PCR Purification Kit (Favorgen-Biotech Corp., Austria), following the manufacturer’s instructions, and subjected to SMRTbell library preparation and PacBio Sequel II sequencing on a single 8M SMRT cell in the University of Oslo, Norway.

### Sequence data workflow

The raw data were obtained in ab1 format for Sanger sequences and fastq format for PacBio sequences. Sanger sequences were inspected for quality using Sequencher v. 5.4.6 software. Sanger sequences were manually checked and trimmed to comprise only the full ITS region or LSU, or a shorter fragment in case of quality issues. PacBio reads were demultiplexed with Lima v.2.4.0 (https://lima.how/; PacBio, 2021) and quality-filtered following Tedersoo et al., (2021). Sequences were trimmed to remove primers using cutadapt v.3.5 (Martin, 2011) and ITS region was extracted using ITSx v.1.1.3 (Bengtsson-Palme et al., 2013). ITS and LSU sequence data were processed separately using seqkit v.2.1.0 (Shen et al., 2016) and VSEARCH v.2.18.0 (Rognes et al., 2016). Reads were grouped into 100% sequence-similarity OTUs. Putative index-switches (also known as tag-jumps) were removed from the OTU table based on the UNCROSS2 score (Edgar, 2018).

The representative reads of each PacBio OTU and all Sanger sequences were identified against UNITE v.9.4b (Nilsson et al., 2018) and SILVA v.138.1 (Quast et al., 2013), respectively, using the BLASTn algorithm and 10 best database hits. Based on morphological identification of specimens, the PacBio OTUs and Sanger sequences were flagged as potentially matching or potentially mismatching to the target taxa. The latter category suggested sequencing of naturally associated fungi, airborne or laboratory contaminants. We did not attempt to distinguish among these groups of unexpected taxa. For a comparison with Sanger sequencing, we only considered non-singleton OTUs with at least 1% relative abundance as putatively true alleles, while others were considered low-quality reads. For each sequenced specimen, we recorded the following properties: the overlap of PacBio OTUs with Sanger sequences from the same specimen (binary), the number of all non-singleton >1%-abundance PacBio OTUs and the number of potentially matching PacBio OTUs.

To calculate pairwise distances within specimens and species, sequences for each sample were aligned using mafft v7.487 (Katoh et al., 2002). Using the dist.seqs command of mothur v1.46.1 (Schloss et al., 2009), we calculated the mean and maximum uncorrected pairwise distances of polymorphic alleles within each sample sequenced for the ITS region. We also recorded corresponding intraspecific distances for 11 species that were successfully sequenced in >5 samples. The samples sequenced for LSU were excluded, because these yielded less data and only a small proportion of samples sequenced for LSU had allele polymorphisms. In allelic comparisons, each pairwise distance represents the percent of mismatches (including indels), where gaps of any length were penalized once and end gaps were ignored.

We also estimated the reasons of sequencing failure. Sanger sequencing was considered failed when the read (1) represented a contaminant; (2) had an unreadable chromatogram; (3) had >5 ambiguous bases; or (4) had >50 bases were missing from either end (due to low-quality 5’ or 3’ end or sequence disruption by length polymorphism of alleles). The PacBio sequencing was considered unsuccessful if the sample (1) yielded no sequence reads; (2) did not contain a correct fungus; or (3) the relative abundance of correct fungus was <10%.

### Statistical analyses

We ran three sets of binomial regression models (logarithmic link function) using the glm function in the R base package. To test whether PacBio sequencing success is more probable for some particular cases of Sanger sequencing failure, we focused on samples that could not be successfully sequenced using Sanger method, and ran a model where PacBio success was a binary dependent variable and Sanger failure reason the sole categorical explanatory variable (four levels, see above). To test if the sequencing success across all samples depended on the relative strength of the PCR band or sample age (years since sampling), we ran models where either Sanger or PacBio success was a binary dependent variable and, respectively, the relative strength of PCR band or sample age was a sole continuous explanatory variable.

## Results

For our samples, the total sequencing costs (incl. library preparation) were ca. 10% less for PacBio than Sanger method. For PacBio, 387 out of 497 samples (77.9%) were successfully sequenced compared with 275 samples when using Sanger (55.3%). Altogether 122 (25%) of all samples could be successfully sequenced with PacBio but not with Sanger, whereas the opposite happened to 15 (3.0%) of samples (Fig 1A). In general, there was a higher probability of successful PacBio outcome in cases where Sanger sequencing failed because of yielding only a partial readable sequence or a sequence containing some low-quality regions (Table 1). The probability for both Sanger and PacBio success increased with the relative strength of the PCR band (Table 1, Fig 2A). The probability of PacBio sequencing success but not Sanger sequencing success significantly decreased with sample age (Table 1, Fig 2B). In 98% of the samples successfully sequenced with PacBio, the true target species was represented by the most abundant OTU.

**Table 1.**
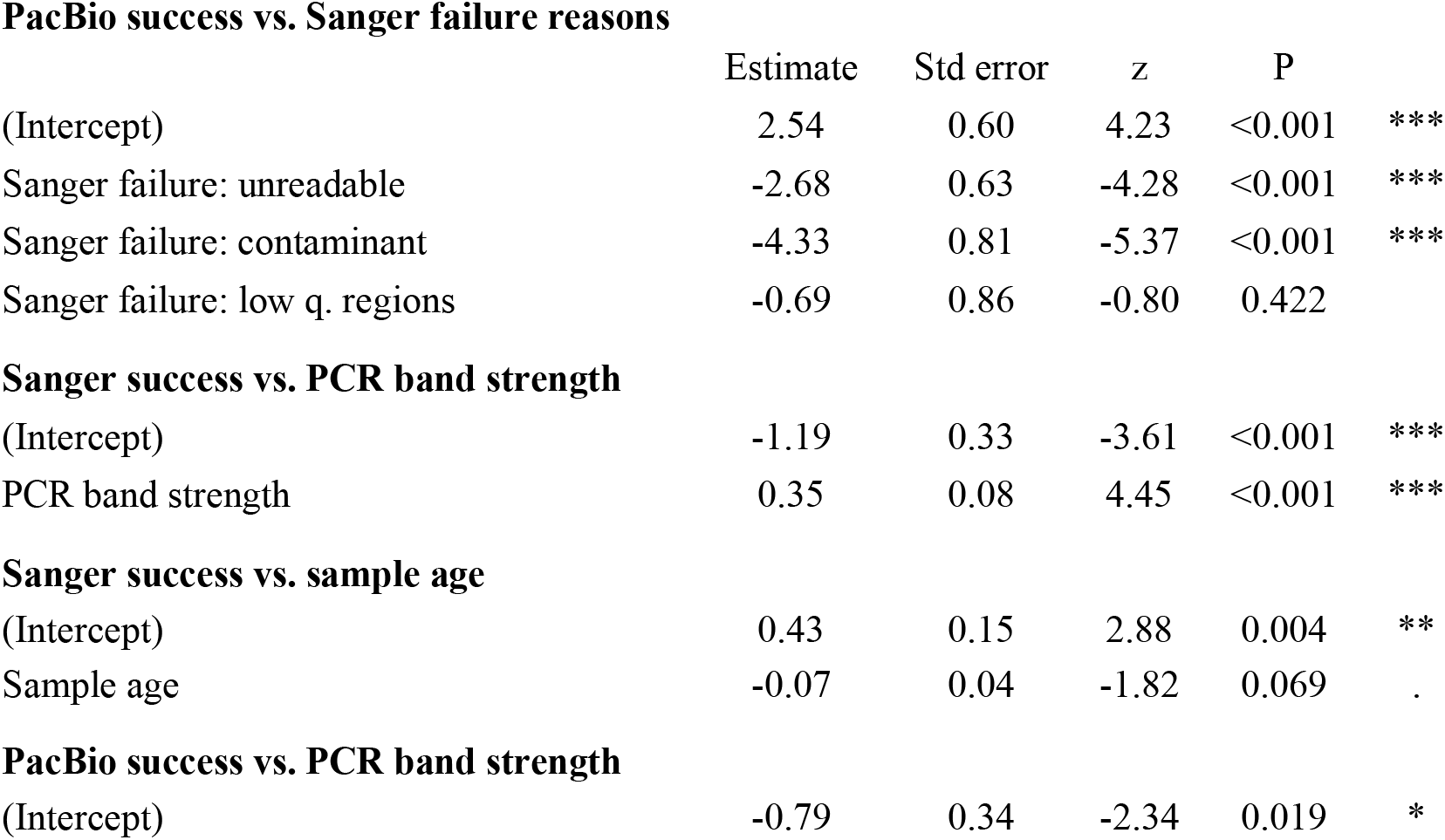
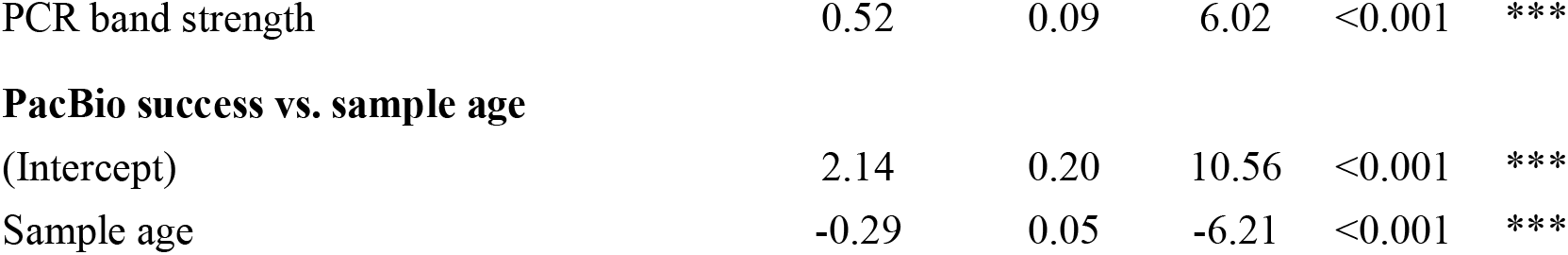
Effects in the binomial regression models. “Sanger failure” is a categorical factor with four levels, where “partial sequence” is a reference group. p-values: * < 0.05; ** < 0.01; *** < 0.001.

**Fig 1.**
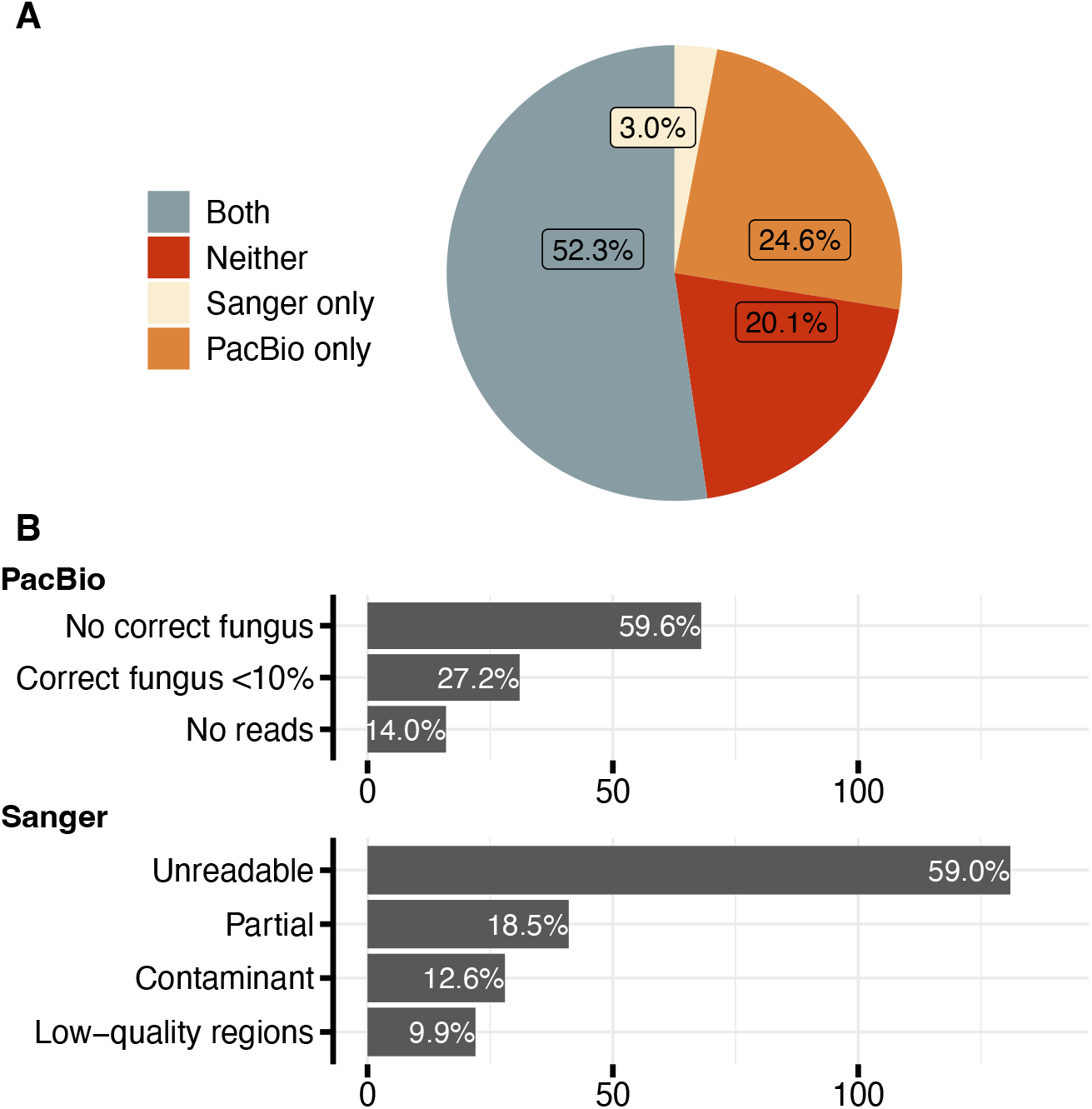
The sequencing success of 497 fungal amplicons with Sanger and PacBio methods (A), and the reasons for failed sequencing (B).

**Fig 2.**
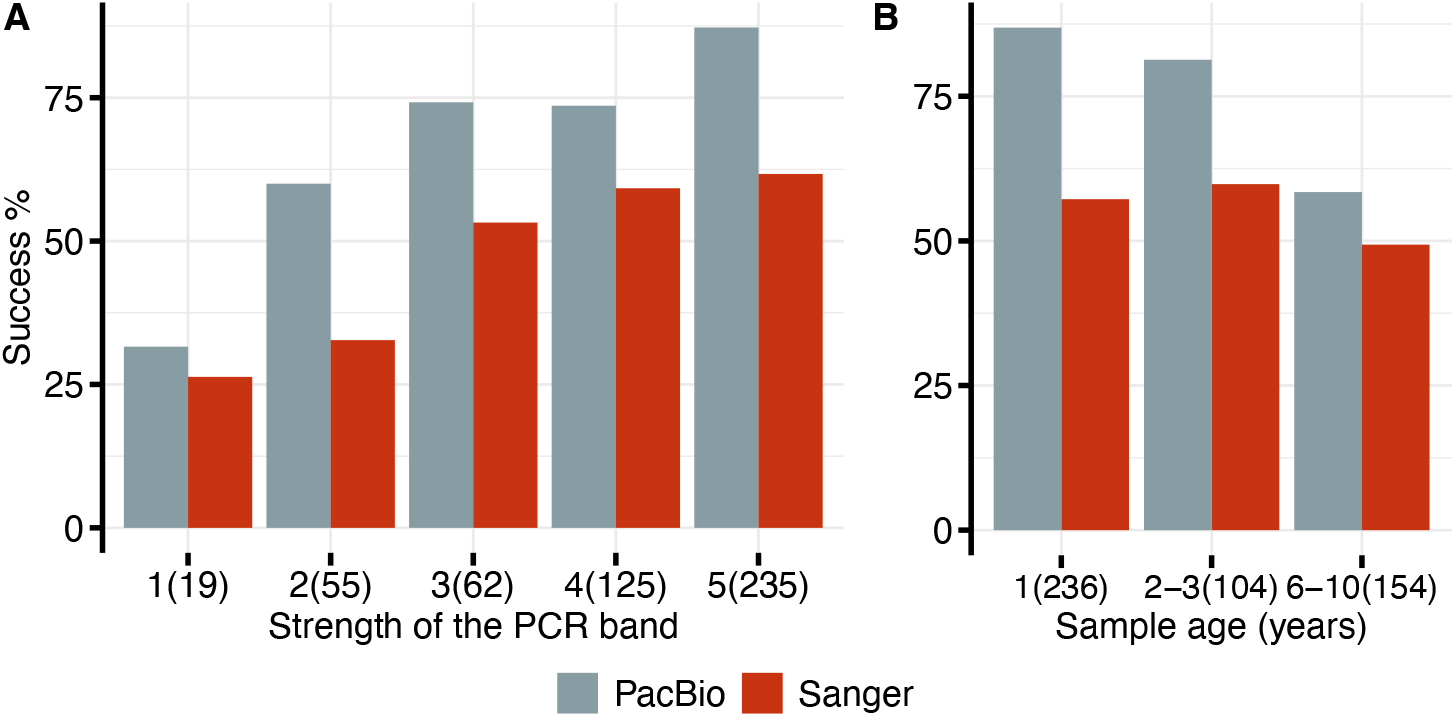
Sequencing success with PacBio and Sanger sequencing at different strengths of the PCR band (A) and sample ages (B). No. of samples in parentheses. There was a single sample with no PCR band.

PacBio sequencing revealed ITS allele polymorphisms in 249 (75%) of samples that were successfully sequenced. These samples contained on average 5.1 (range 2 to16) polymorphic alleles with >1% relative abundance. Each allele generally differed from all others by only a few base pairs, yielding an average intraindividual distance of 0.44% (range, 0.15% to 2.88%) across all samples with allele polymorphisms. The average maximum distance across all samples with allele polymorphisms was 0.64% (range, 0.15% to 2.88%) (Fig 3B). The relative abundance of the most abundant allele and the total number of sequences in the sample were weakly correlated (Spearman r = 0.13, Fig 3A).

**Fig 3.**
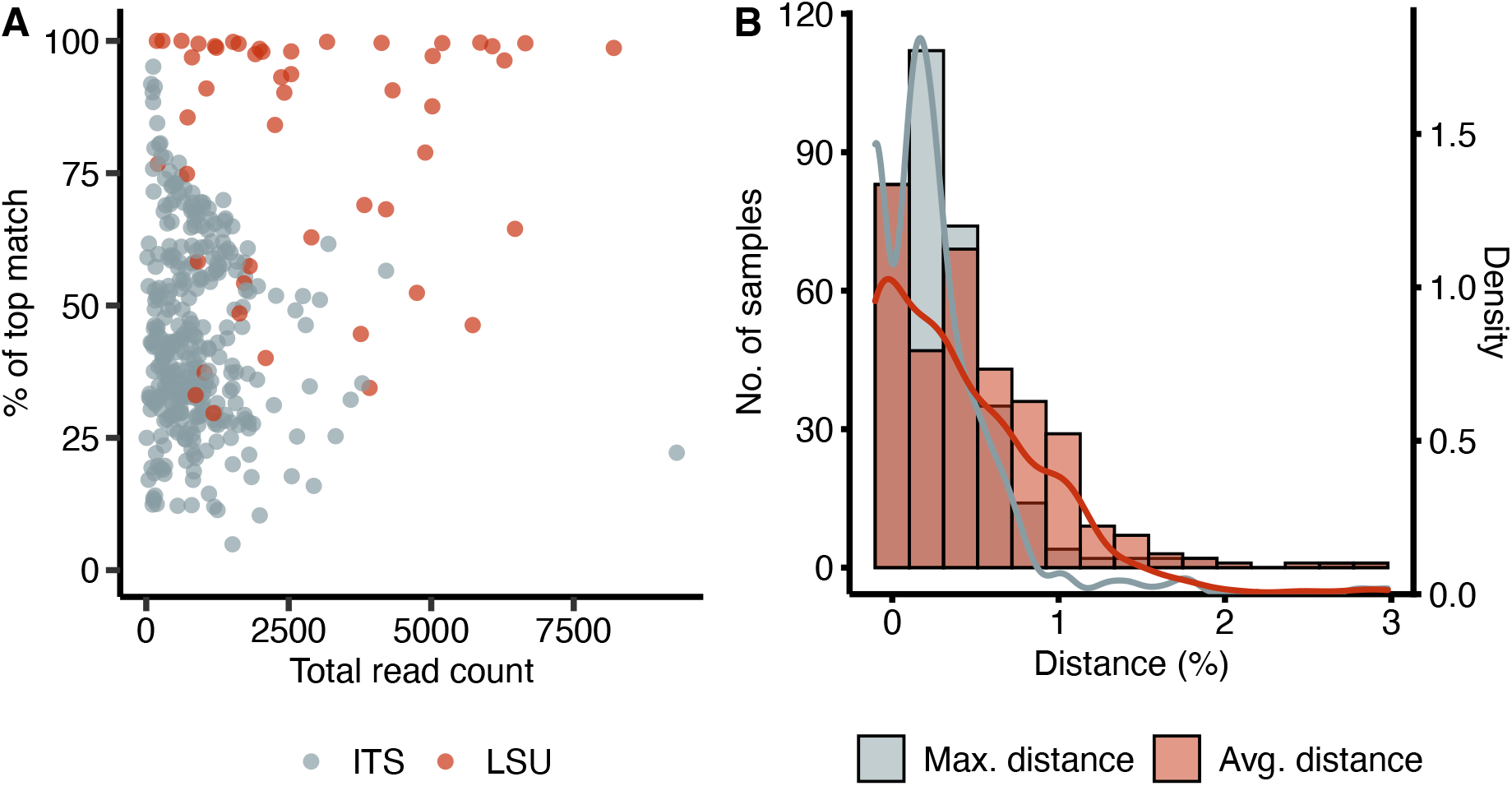
The major characteristics of successful PacBio samples. A: The relationships between total no. of sequences and relative abundance of the top (most abundant) allele. B: Count and density (probability density) of average and maximum distances between alleles in samples successfully sequenced for the ITS region. Distance is zero for samples with no allele polymorphism.

We also checked among-individual, intraspecific variation for 11 species that were successfully sequenced in >5 samples (Table 2). Those species had in average 2.5 polymorphic alleles per sequenced sample (min 0.29, max 5.17). The average and maximum intraspecific distance remained below 2% in all cases. The polypore *Sidera vulgaris* displayed the highest average and maximum intraspecific differences.

**Table 2.** Number of polymorphic alleles, and intraspecific distances in 11 studied species.

|  | No. of<br>samples | No. of<br>polymorphic<br>alleles | Min<br>distance<br>(%) | Max<br>distance<br>(%) | Average<br>distance<br>(%) |
| --- | --- | --- | --- | --- | --- |
| *Antrodia piceata* | 8 | 6 | 0.171 | 0.512 | 0.262 |
| *Antrodia serialis* | 6 | 19 | 0.181 | 1.093 | 0.565 |
| *Physisporinus vitreus* L4 | 7 | 9 | 0.183 | 0.730 | 0.386 |
| *Postia tephroleuca* | 6 | 12 | 0.185 | 0.750 | 0.431 |
| *Rhodonia placenta* | 10 | 24 | 0.172 | 1.382 | 0.717 |
| *Sidera vulgaris* | 33 | 140 | 0.181 | 1.828 | 0.911 |
| *Skeletocutis nemoralis* | 10 | 35 | 0.167 | 1.003 | 0.510 |
| *Skeletocutis semipileata* | 14 | 61 | 0.166 | 1.815 | 0.720 |
| *Skeletocutis stellae* | 6 | 31 | 0.174 | 1.394 | 0.681 |
| *Tomentella* sp. 15 | 6 | 2 | 0.173 | 0.173 | 0.173 |
| *Tomentella* sp. 23 | 7 | 2 | 0.175 | 0.175 | 0.175 |

## Discussion

### 1. Sequencing success

We demonstrated that sequencing of nearly 500 fungal amplicons using long-read HTS sequencing (PacBio) had a higher success rate compared to the traditional Sanger sequencing at comparable cost. Identifying the target species from PacBio samples was usually straightforward, as these were represented by the dominant OTU. However, contaminants were common and sometimes prevailed, especially in relatively old specimens. This indicates that taxonomic knowledge remains important when interpreting sequencing results (see Vu et al., 2018). We observed a rapid decay in PacBio sequencing success rate in >5-year-old specimens. Earlier studies have noticed a similar trend in Sanger sequencing (Larsson & Jacobsson, 2004). In the case of Sanger, we used single-end sequencing (as most taxonomic studies), additional sequencing of the complementary strand would have increased the success rate in samples with partial sequences, and low quality regions (Hyde et al., 2013).

There were two typical situations where the Sanger sequencing failed, but PacBio proved successful: length polymorphism of reads and field-contaminated samples. The length polymorphism in alleles caused disruption of Sanger reads but yielded no issues in PacBio. For example, in the boreal polypore species *Sidera vulgaris*, Sanger sequencing recovered a full-length read in 7% of the sequenced specimens, whereas PacBio was successful in 97% of specimens. The field-contamination was common in collections of the resupinate tropical *Tomentella* spp. and polypore *Rhodonia placenta*. In *R. placenta*, Sanger sequencing and PacBio sequencing were successful in 27% and 91% out of 11 specimens analyzed, respectively. It is worth noticing that we also adopted relatively stringent criteria for successful sequencing. If fruit body characteristics allowed restricting the species identification to a few options only, many of the “ sequencing failures” were still informative for confirming the final identification.

### 2. Allele polymorphism in studied samples

Our workflow pointed to a widespread allele polymorphism within the ITS marker in the studied fungal collections. We consider most of the common variants as “ true” alleles, since PacBio circular consensus sequencing yields high sequencing accuracy (Karst et al., 2021; Tedersoo et al., 2021) and we only addressed OTUs represented by >1 read and at least 1% total abundance, hence avoiding random PCR and sequencing errors (see also Ganley & Kobayashi, 2007). Our interpretation contrasts to Lindner et al., (2013), who ascribed intraindividual variation in a majority of 100 sampled fungal species to PCR and sequencing errors that were an order of magnitude more common in the now-obsolete 454 pyrosequencing technology (Lindner et al., 2013). However, the alleles typically differed by a single or few positions, and the maximum intragenomic and intraspecific differences remained <2% (except in three samples). Metabarcoding studies typically use a 97%, 98% or 98.5% ITS sequence similarity threshold for species-level separation and identification of taxa. Therefore, the observed differences typically remain within this threshold, especially when compared to the closest read (single-linkage clustering) or centroid (greedy clustering) based methods. However, when combined with geographic distance (population divergence), inappropriate clustering methods and accumulating sequencing errors, intraspecific differences may indeed account for artefactual, elevated richness in ecological studies.

Our results suggest that the exact sequence variant (ESV) based approaches (Callahan et al., 2017) are not optimal for species-level metabarcoding analyses of fungal diversity (see also Estensmo et al., 2021; Tedersoo et al., 2022) and perhaps eukaryotes in general (Antich et al., 2021; Porter & Hajibabaei, 2021), by potentially retrieving artefactual taxa. In conclusion, multiplex DNA barcoding of the fungal ITS marker using a PacBio third-generation HTS protocol is a useful tool for taxonomic assessment of large sets of vouchered fungal specimens. Besides costs comparable to Sanger sequencing, PacBio HTS provides more complete and accurate recovery of various alleles, which can potentially be accounted for in bordering the molecular species or species hypotheses (Kõljalg et al., 2013) and used in population-level studies (Byrne et al., 2017).

## Acknowledgements

The study was funded by the State Forest Management Centre (project “ Enhancing the conservation performance of protected forest fragments” to K.R.), the Estonian Research Council (grants PRG632 and PRG1170), and the European Regional Development Fund (Centre of Excellence EcolChange). Rasmus Puusepp performed the laboratory work.

## Data accessibility and benefit-sharing

Data accessibility: After the paper is published, the unique haplotype data will be made available through the PlutoF web platform.

Benefits Generated: Benefits from this research accrue from the sharing of our data and results on public databases as described above.

## Author contributions

L.T and K.R conceived the research idea and designed the study, K.R and U.K collected data, K.R, L.T, I.S, V.M and O.C analyzed data, K.R and L.T wrote the paper, all authors discussed the results and commented on the manuscript.

